# Complement and coagulation cascades are potentially involved in dopaminergic neurodegeneration in α-synuclein-based mouse models of Parkinson’s disease

**DOI:** 10.1101/2020.01.11.900886

**Authors:** Shi-Xun Ma, Donghoon Kim, Yulan Xiong, Seung-Hwan Kwon, Saurav Brahmachari, Sangjune Kim, Tae-In Kam, Raja Sekhar Nirujogi, Sang Ho Kwon, Valina L. Dawson, Ted M. Dawson, Akhilesh Pandey, Chan Hyun Na, Han Seok Ko

## Abstract

Parkinson’s disease (PD) is the second most common neurodegenerative disorder that results in motor dysfunction and eventually, cognitive impairment. α-Synuclein protein has been known to be the most culprit protein, but the underlying pathological mechanism still remains to be elucidated. As an effort to clarify the pathogenesis mechanism by α-synuclein, various PD mouse models with α-synuclein overexpression have been developed. However, the systemic analysis of protein abundance change by the overexpressed α-synuclein in the whole proteome level has been still lacking. To address this issue, we established two sophisticated mouse models of PD by injecting α-synuclein preformed fibrils (PFF) or by inducing overexpression of human A53T α-synuclein to discover overlapping pathways, which could be altered in the two different types of PD mouse model. For more accurate quantification of mouse brain proteome, stable isotope labeling with amino acid in mammal-based quantification was implemented. As a result, we have successfully identified a total of 8,355 proteins from both of the mouse models; ∼6,800 and ∼7,200 proteins from α-synuclein PFF injected mice and human A53T α-synuclein transgenic mice, respectively. From the pathway analysis of the differentially expressed proteins in common, the complement and coagulation cascade pathway were determined as the most enriched ones. This is the first study that highlights the significance of the complement and coagulation pathway in the pathogenesis of PD through proteome analyses with two sophisticated mouse models of PD.

## Introduction

Parkinson’s disease (PD) is the second most prevalent neurodegenerative disease (1). Although the pathological mechanism has not been well-defined, numerous culprit genes such as *SNCA, LRRK2, Parkin, PINK1, DJ-1* and *ATP13A2* have been reported (2). In detail, *SCNA* and *LRRK2* are tightly linked to autosomal-dominant PD forms, whereas *Parkin, PINK1, DJ-1* and *ATP13A2* are majorly implicated in autosomal-recessive PD forms (3). Among these culprit genes, *SNCA* is the first identified gene that serves as a major cause of PD development when it is mutated (4). Since the human α-synuclein is encoded by the *SNCA*, researchers have focused on the functional role of α-synuclein in PD. In the past few years, many studies have demonstrated that the formation of abnormal aggregation of α-synuclein, which we now call Lewy bodies (LB) and Lewy neurites, is one of the pathological hallmarks of the PD (5).

For a more-in-depth comprehension of the underlying mechanisms of PD, several animal models such as toxin-based models, α-synuclein genetic models, and α-synuclein-aggregates injected models have been generated (6–8). The neurotoxins, such as 1-methyl-4-phenyl-1,2,3,6-tetrahydropyridine (MPTP), 6-hydroxydopamine (6-OHDA), and rotenone, are widely used to produce the toxin-based PD model. Although these neurotoxins induce the vast and rapid degeneration of the nigrostriatal pathway, all of the neurotoxin-based models essentially lack the representative PD hallmark, LB (9). Currently, various α-synuclein transgenic animal models have been developed using different promoters for driving the expression of α-synuclein transgene codes of wild type or the mutants such as A53T, A30P, or E46K (6). In some of these models, decreased striatal levels of tyrosine hydroxylase (TH) or dopamine (DA) along with the resulting motor behavioral deficits were observed, indicating that the accumulation of α-synuclein can significantly alter the functioning of DA neurons. Obtrusive nigrostriatal degeneration, however, has been rarely detected in most of the animal models. Indeed, the prion protein promoter driving A53T α-synuclein transgenic mouse model failed to reproduce the DA neuronal loss in the substantia nigra par compacta (SNpc) or brainstem (10). Interestingly, it has been recently reported that a new line of mouse model in which α-synuclein expression is dictated under the control of the paired like homeodomain 3 promoter in the DA neurons with tetracycline-regulated inducible system showed exemplar dopaminergic neuronal degeneration, motor deficits, decreasing of DA release, and impairments of the autophagy-degradation pathway (11). To date, several studies have demonstrated prion-like cell-to-cell transmission of pathological α-synuclein aggregates. Embodying this remarkable pathological feature in an animal model, Luk *et al.* created a PD-like mouse model by a single intrastriatal injection of synthetic α-synuclein pre-formed fibrils (PFF), which in turn allows for cell-to-cell transmission of pathological α-synuclein aggregates, formation of LB pathology, progressive DA neuronal degeneration in the SNpc, and motor behavioral deficits (8).

The proteomic findings from animal models of PD, such as MPTP-treated mouse model, 6-OHDA-injected rat model, and A30P*A53T α-synuclein transgenic mouse models, provided many valuable insights into several pathways that may be related to disease pathogenesis (12–15). However, those proteomic analyses still showed limitations as illustrated by their shallow proteome analysis or analysis of PD model mouse brains without visible PD-like phenotypes. Notably, these studies were all conducted with a single PD mouse model. Although PD mouse models are generated to recapitulate PD pathogenesis that happens in the human brain, proteins differentially expressed in those mouse models are a mixture of artifact proteins and *bona fide* proteins involved PD pathogenesis. For this reason, we postulated that the use of multiple PD models would provide an advantage over using a single mouse model for exploring the possibilities for the most pronounced pathway involved in PD pathogenesis that happens in the human brain. For this reason, we generated two different types of PD model mice by stereotaxic injections of α-synuclein PFF into the striatum of wild type mice or adeno-associated virus (AAV)-tetracycline-regulated transactivator (tTA) into tetracycline promoter (TetP)/A53T α-synuclein mice, followed by thorough characterization of PD phenotypes to ensure the occurrence of pathological phenotype.

The mass spectrometry-based proteomics has been widely used for global protein quantification from various biological samples. Multiple protein quantification methods using the mass spectrometry technique has been developed. Because the most commonly used and the most accurate quantification method is a metabolic labeling-based quantification (16, 17), we took stable isotope labeling with amino acid in mammals (SILAM) approach where the relative protein abundance is quantified by spiking in mouse samples completely labeled with heavy lysine to the samples to be measured. In order to discover affected pathways in both PD model mice, a pathway analysis was also conducted.

This is the first proteomics study in which two different types of PD mouse model with key phenotypes observed in PD patients are analyzed to search for the affected pathways and thus explain α-synuclein induced neurodegeneration. This study would pave the way to the elucidation of the PD pathogenesis.

## Experimental Procedures

### Animal models

All experimental procedures were followed according to the guidelines of Laboratory Animal Manual of the National Institute of Health Guide to the Care and Use of Animals, which were approved by the Johns Hopkins Medical Institute Animal Care and Use Committee. (1) Mouse strain for stereotaxic α-synuclein PFFs injection. C57BL6 mice were obtained from the Jackson Laboratories (ME, USA). (2) Conditional TetP-hA53T α-synuclein transgenic mouse. To generate pPrP-TetP-hA53T α-synuclein mice, the cDNA encoding human WT α-synuclein was subcloned into the unique XhoI site of the 9.0 kb pPrP-TetP vector. Site directed mutagenesis was conducted with the pPrP-TetP-WT α-synuclein construct as a template to generate pPrP-TetP-A53T α-synuclein construct. hA53T mutation was confirmed by DNA sequencing. The pPrP-TetP-A53T α-synuclein construct was linearized with digestion with Not1 and the purified linearized DNA fragment (7 kb) was used for pronuclear microinjection of single-cell embryos from B6C3F2 strain and the one or two cell embryos were transferred into B6D2F1 pseudo-pregnant female mice to drive founder mice. This was conducted by the National Cancer Institute Transgenic Core Facility. Founder animals were screened for transgene incorporation by PCR of tail genomic DNA using TetP-α-synuclein primers (Forward: CGG GTC GAG TAG GCG TGT AC; Reverse: TCT AGA TGA TCC CCG GGT ACC GAG: PCR product: 173 bp). Positive founder mice with a high copy number of the transgene were crossed to B6 mice. A TetP-hA53T α-synuclein mouse line was generated in our previous study (18).

### Generation of SILAM mouse and validation of the labeling efficiency

F2 SILAM wide type female mice were purchased from Cambridge Isotope Laboratories and then crossed with regular C57BL6 wild type males by fed with stable isotope labeling using amino acids in cell culture (SILAC) L-lysine (^13^C_6_) feed (Cambridge Isotope Laboratories, Inc). After 7 days, the males were separated. F3 SILAM pups were weaned at 28 days old. At two months of age, F3 SILAM mice were anesthetized by isoflurane, euthanized by decapitation followed by harvesting tissue samples. During the entire period, the mice were fed with SILAC L-Lysine (^13^C_6_) feed purchased from Cambridge Isotope Laboratories. Heavy label incorporation of L-Lysine (^13^C_6_) was performed by mass spectrometry. Briefly, the dissected brain sample was solubilized in the lysis buffer (8M Urea in 50mM TEABC pH8.0 containing protease inhibitors) and sonicated using Branson sonicator probe to shear DNA. The lysate was spun down at 20,000 xg for 20 min and the supernatant was collected and the protein amount was determined using BCA assay. 20 μg of the lysate was reduced with 5mM DTT by incubating at room temperature for 1 hr and alkylated with 20mM IAA by incubating at room temperature in the dark. Further, the lysate was subjected to trypsin digestion at 1:20 ratio and incubated at 37°C overnight. The same procedure was followed for the serum sample. Tryptic peptides were acidified and desalted using C18 stage-tips and eluted peptides were vacuum dried. The prepared peptides were analyzed on Orbitrap Elite Mass spectrometer that is coupled with Easy nanoLC II. Peptides were resolved on a homemade 15 cm column (C18, 3μm, 100 A, magic C18). A linear gradient of 3 to 35% was applied to resolve the peptides for about 35 minutes and a total run time of 45 minutes. The mass spectrometer was operated in a data-dependent mode by targeting top 10 peptide precursor ions. Precursor ions were measured at 120,000 resolution at m/z 400 using Orbitrap mass analyzer and the produced ions were fragmented using HCD at 32% normalized collision energy and measured at 15,000 resolution at m/z 400 using Orbitrap mass analyzer. The raw mass spectrometry data was searched using Proteome Discoverer 1.4 software suite using Sequest search algorithm against a Refseq 58 mouse protein database. To determine the heavy label incorporation efficiency the SILAC Lys 13C6 modification was set as variable modification along with the Oxidation of Met. Carbamidomethylation of Cys was set as a fixed modification. 1% peptide and PSM FDR was applied and rank 1 peptides were filtered and retained for the final analysis.

### α-synuclein PFF preparation

Recombinant mouse α-synuclein proteins were purified as previously described with IPTG independent inducible pRK172 vector system (19–23). Briefly, bacteria were harvested by centrifugation at 6,000 x g for 10 min after 16 h incubation at 37C. The bacterial pellet was resuspended in high salt buffer containing 10 mM Tris (pH 7.6), 750 mM NaCl, and 1 mM EDTA with complete protease inhibitor mixture (Sigma-Aldrich) and 1 mM PMSF, and lysed by sonicating for 5 min (30 s pulse on/off) at 60% amplitude (Branson Digital sonifier, Danbury, CT, USA) with boiling for 15 min. After centrifugation at 6,000 x g for 20 min, the supernatant was subjected to serial purification steps using Superdex 200 Increase 10/300 G size-exclusion and Hitrap Q Sepharose Fast Flow anion-exchange columns (GE Healthcare, Pittsburgh, PA, USA). Purified a-syn was applied to High capacity endotoxin removal spin columns (Pierce, Rockford, IL, USA) and Ni Sepharose 6 Fast Flow (GE Healthcare) to remove endotoxin, followed by confirmation of removal of endotoxin using LAL Chromogenic Endotoxin Quantitation Kit (Pierce). α-synuclein monomer was stored at −80 until used. α-synuclein PFF was prepared in PBS from 5 mg/ml of α-synuclein monomer by stirring with magnetic stirrer (1,000 rpm at 37 °C) for 1 week. α-synuclein aggregates were sonicated for 30 s (0.5 s pulse on/off) at 10% amplitude (Branson Digital sonifier).

### Stereotaxic α-synuclein PFFs and AAV1-tTA-virus injection

For stereotaxic injection of α-synuclein PFFs and adeno-associated virus 1 (AAV1)-tetracycline-regulated transactivator (tTA), three-month-old male and female mice were anesthetized with xylazine and ketamine. (**1**) An injection cannula (26.5 gauge) was applied stereotaxically into the striatum (mediolateral, 2.0 mm from bregma; anteroposterior, 0.2 mm; dorsoventral, 2.6 mm) unilaterally (applied into the right hemisphere). The infusion was performed at a rate of 0.2 μl per min, and 2 μl of α-synuclein PFFs (2.5 μg/μl in PBS) or same volume PBS were injected into the mouse. (**2**) An injection cannula (26.5 gauge) was applied stereotaxically into the SNpc (anteroposterior, 3.2 mm from bregma; mediolateral, 1.3 mm; dorsoventral, 4.3 mm) unilaterally (applied into the right hemisphere). The infusion was performed at a rate of 0.2 μl per min, and 2 or 1 μl of a high-titer AAV1-tTA-GFP (3.5 × 10^13^ AAV vector genomes per ml in PBS) and AAV1-GFP was injected into PrP-TetP-hA53T α-synuclein mice. After the final injection, the injection cannula was maintained in the striatum and substantia nigra for an additional 5 min for complete absorption of the PFFs and virus and then slowly removed from the mouse brain. The head skin was closed by suturing, and wound healing and recovery were monitored following surgery.

### Immunohistochemistry and quantitative analysis

Immunohistochemistry (IHC) was performed on 30 μm thick serial brain sections. Primary antibodies and working dilutions are detailed in Supplemental Table S1. For histological studies, perfusion was performed with ice-cold phosphate buffered saline (PBS) and the brains were fixed with 4% paraformaldehyde/PBS (pH 7.4). The brains were collected and post fixed for 4 h in 4% paraformaldehyde followed by cryoprotection in 30% sucrose/PBS (pH 7.4) solution. Brains were frozen in OCT buffer and serial coronal sections (30 μm sections) were cut with a microtome. Free-floating 30 μm sections were blocked with 4% goat or horse serum/PBS plus 0.2% Triton X-100 and incubated with an antibody against TH (Novus Biologicals, Littleton, CO, USA) followed by incubation with biotin-conjugated anti-rabbit antibody or biotin-conjugated anti-mouse antibody (Vectastain Elite ABC Kit, Vector Laboratories, Burlingame, CA, USA). After triple washing steps, ABC reagents (Vectastain Elite ABC Kit, Vector Laboratories) were added and sections were developed using SigmaFast DAB Peroxidase Substrate (Sigma-Aldrich). Sections were counterstained with Nissl (0.09% thionin). TH-positive and Nissl positive DA neurons from the SNpc region were counted through optical fractionators, the unbiased method for cell counting. This method was carried out by using a computer-assisted image analysis system consisting of an Axiophot photomicroscope (Carl Zeiss Vision) equipped with a computer controlled motorized stage (Ludl Electronics), a Hitachi HV C20 camera, and Stereo Investigator software (MicroBright-Field). The total number of TH-stained neurons and Nissl counts were analyzed as previously described (24). Fiber density in the striatum was analyzed by optical density (OD) measurement. ImageJ software (NIH, http://rsb.info.nih.gov/ij/) was used to analyze the OD as previously described (24, 25).

### Immunofluorescence analysis

Immunofluorescence was performed on 30 μm thick serial brain sections. Primary antibodies and working dilutions are detailed in Supplementary Table S1. For histological studies, immunofluorescence in tissue sections was performed as described previously with some modifications (19). Paraformaldehyde (4% in PBS, pH 7.4)-fixed coronal brain sections were blocked with 10% donkey serum (Jackson Immunoresearch)/PBS plus 0.3% Triton X-100 in PBS and incubated with antibodies to p-S^129^-α-synuclein (1:1000; Biolegend), and TH (1:1000; Novus Biologicals), for overnight at 4° C. After brief washes with PBS, floating brain sections were incubated with 0.1% Triton X-100 and 5% donkey serum in PBS, followed by 1 hr of incubation with a mixture of FITC-conjugated (Jackson Immunoresearch) and CY3-conjugated (Jackson Immunoresearch) secondary antibodies at room temperature. The fluorescent images were acquired via a Zeiss confocal microscope (Zeiss Confocal LSM 710) after the coverslips were mounted with DAPI mounting solution (VECTASHIELD HardSet Antifade Mounting Medium with DAPI, Vector laboratories). All images were processed by the Zeiss Zen software. The selected area in the signal intensity range of the threshold was measured using ImageJ analysis.

### Immunoblot analysis

Mouse brain tissues were homogenized and prepared in lysis buffer [10 mM Tris-HCL, pH 7.4, 150 mM NaCl, 5 mM EDTA, 0.5% Nonidet P-40, 10 mM Na-β-glycerophosphate, phosphate inhibitor mixture I and II (Sigma-Aldrich, St. Louis, MO, USA), and complete protease inhibitor mixture (Roche, Indianapolis, IN, USA)], using a Diax 900 homogenizer (Sigma-Aldrich). After homogenization, samples were rotated at 4° C for 30 min for complete lysis, the homogenate was centrifuged at 22,000 x g for 20 min and the supernatants were collected. Protein levels were quantified using the BCA Kit (Pierce, Rockford, IL, USA) with BSA standards and analyzed by immunoblot. Electrophoresis on 8–16% and 4–20% gradient SDS-PAGE gels was performed in order to resolve the proteins (obtained 10-20 μg) from the mouse brain tissue or cell lysates. The proteins were then transferred to nitrocellulose membranes. The membranes were blocked with blocking solution (Tris-buffered saline with 5% non-fat dry milk and 0.1% Tween-20) for 1 hr and incubated at 4° C overnight with anti-TH (Novus Biologicals), anti-Factor H (Abcam), anti-C3 (Abcam), anti-Fibrinogen (Abcam) and anti-α-synuclein (BD bioscience) antibodies, followed by HRP-conjugated goat of mouse secondary antibodies (Jackson ImmunoResearch), HRP-conjugated rabbit of sheep secondary antibodies (1:3000 ThermoFisher) and HRP-conjugated goat of rabbit secondary antibodies (1: 3000, Jackson ImmunoResearch) for 1 hr at room temperature. Primary antibodies and working dilutions are detailed in Table xx. The bands were visualized by enhanced chemiluminescence (Thermo Scientific, IL, USA). Finally, the membranes were re-probed with HRP-conjugated β-actin antibody (Sigma-Aldrich) after it was stripped.

**Table 1.**
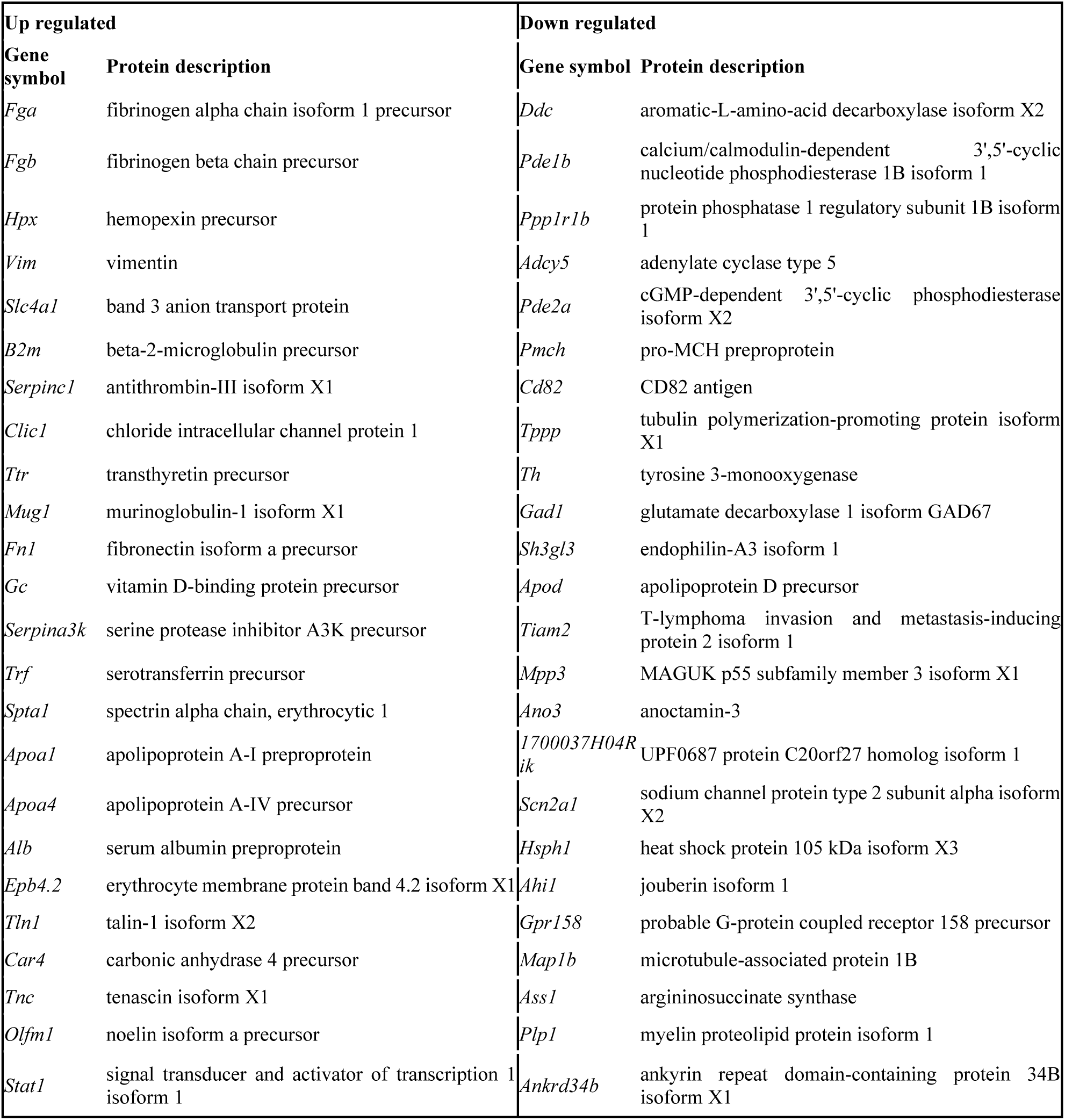
List of differential proteins common in both α-synuclein PFFs injected mice and hA53T α-synuclein Tg mice

### Behavioral tests

The experimenter was blinded to the treatment group for all behavioral studies. All tests were recorded and performed between 10:00-16:00 in the lights-on cycle. (1) **Pole test.** Mice were acclimatized in the behavioral procedure room for 30 min. The pole was made of a 75 cm metal rod with a diameter of 9 mm. It was wrapped with bandage gauze. Mice were placed on the top of the pole (7.5 cm from the top of the pole) facing head-up. The total time taken to reach the base of the pole was recorded. Before the actual test, mice were trained for two consecutive days. Each training session consisted of three test trials. On the test day, mice were evaluated in three sessions and the total time was recorded. The maximum cutoff time to stop the test and recording was 60 sec. Results for turn down, climb down, and total time (in sec) were recorded. (2) **Rotarod test.** For the rotarod test, mice were placed on an accelerating rotarod cylinder, and the time the animals remained on the rotarod was measured. The speed was slowly increased from 4 to 40 rpm within 5 min. A trial ended if the animal fell off the rungs or gripped the device and spun around for 2 consecutive revolutions without attempting to walk on the rungs. The animals were trained 3 days before the test. Motor test data are presented as the percentage of the mean duration (3 trials) on the rotarod compared with the control.

### Sample preparation for proteome analysis of PD model mouse samples

The mouse brain samples were extracted by Filter Aided Sample Prep (FASP) approach (26). Briefly, the mouse ventral midbrain samples were sonicated in 50 mM triethylammonium bicarbonate (TEAB)/4% SDS/10 mM dithiothreitol with Halt protease inhibitor cocktail (Thermo Scientific) for 5 min followed by heating at 95°C for 5 min. After cooling down, the proteins were alkylated with 30 mM iodoacetamide at room temperature for 30 min followed by centrifugation at 16,000 x g for 10 min. SDS in the sample was removed by diluting the samples with 50 mM TEAB/8 M urea and concentrated with a centricon with 30 kDa MWCO for 40 min at RT. This buffer exchange step was repeated five times. Sweat gland proteins were digested with Lys-C at room temperature for 3 h followed by further digestion with trypsin overnight. Following enzyme digestion by either approach, the peptides were desalted using Sep-Pak C_18_ cartridge and fractionated into 24 fractions by basic pH reversed-phase liquid chromatography. Briefly, lyophilized samples were reconstituted in solvent A (10 mM TEAB, pH 8.5) and loaded onto XBridge C_18_, 5 μm 250 × 4.6 mm column (Waters, Milford, MA). Peptides were resolved using a gradient of 3 to 50% solvent B (10 mM TEAB in acetonitrile, pH 8.5) over 50 min collecting 96 fractions. The fractions were subsequently concatenated into 24 fractions followed by vacuum drying using SpeedVac. The dried peptides were reconstituted in 15 μl 10% FA and the entire amount was injected.

### Mass spectrometric analysis of PD model mouse samples

The fractionated peptides were analyzed on an LTQ-Orbitrap Elite mass spectrometer (Thermo Scientific, Bremen, Germany) coupled to EASY-nanoLC II system. The peptides from each fraction were reconstituted in 10% formic acid and loaded onto a trap column (100 μm x 2 cm) at a flow rate of 5 μl per minute. The loaded peptides were resolved at 250 nl/min flow rate using a linear gradient of 10 to 35% solvent B (0.1% formic acid in 95% acetonitrile) over 95 minutes on an analytical column (50 cm x 75 μm ID) packed in house and was fitted on EASY-Spray ion source that was operated at 2.0 kV voltage. Mass spectrometry analysis was carried out in a data dependent manner with a full scan in the range of *m/z* 350 to 1550 in top speed mode setting 3 sec per cycle. Both MS and MS/MS were acquired and measured using Orbitrap mass analyzer. Full MS scans were measured at a resolution of 120,000 at *m/z* 400. Precursor ions were fragmented using higher-energy collisional dissociation (HCD) method and detected at a mass resolution of 30,000 at *m/z* 400. The HCD energy was set to 32 for mouse samples and 35 for human samples. The Automatic gain control for full MS was set to 1 million ions and for MS/MS was set to 0.05 million ions with a maximum ion injection time of 50 and 100 ms, respectively. Dynamic exclusion was set to 30 sec and singly charged ions were rejected. Internal calibration was carried out using a lock mass option (*m/z* 445.1200025) from ambient air.

### Data analysis

Proteome Discoverer (v 2.1; Thermo Scientific) suite was used for identification and quantitation. The tandem mass spectrometry data were searched using SEQUEST search algorithms against a human RefSeq database (version 70) supplemented with frequently observed contaminants. The search parameters used were as follows: a) trypsin as a proteolytic enzyme (with up to two missed cleavages) b) peptide mass error tolerance of 10 ppm; c) fragment mass error tolerance of 0.02 Da; d) Carbamidomethylation of cysteine (+57.02146 Da) as fixed modification and oxidation of methionine (+15.99492 Da) as variable modifications. The minimum peptide length was set to 6 amino acids. Peptides and proteins were filtered at a 1 % false-discovery rate (FDR) at the PSM level using percolator node and at the protein level using protein FDR validator node, respectively.

### Pathway analysis

Pathway analysis Pathway analysis was performed at DAVID.(27, 28) All mass spectrometry data and search results have been deposited to the ProteomeXchange Consortium (http://proteomecentral.proteomexchange.org/cgi/GetDataset?ID=PXD015293) via the PRIDE partner repository with the dataset identifier PXD015293 and project name ‘Quantitative proteome analyses of two types of Parkinson’s disease mouse model revealed potential pathways of the pathogenesis’. Reviewers can access the dataset by using ‘’ as ID and ‘e845DhoI’ as password.

### Statistics

For the validation of mouse phenotype, data were presented as mean ± s.e.m from at least three independent experiments. In order to assess the statistical significance, Student’s t-test or ANOVA tests followed by Bonferroni post hoc test were performed using Prism 6 software (GraphPad). Assessments with a p < 0.05 were considered significant. The proteome analysis has been conducted in technical duplicate. The P values of the mass spectrometry data were calculated by the Student’s two-sample t-test. The fold changes were calculated by dividing the average abundance of one group by the one of another group. The q-values were calculated by Significance Analysis of Microarrays and a permutation-based false discovery rate estimation to avoid the identification of false positive differential proteins that happens when multiple comparisons are tested (29).

## Results

To identify pathways commonly, yet critically involved in PD pathogenesis, we generated two different types of PD mouse model by injecting α-synuclein PFFs into the striatum of wild type mice or by injecting AAV1-tTA into the ventral midbrain of TetP-A53T α-synuclein mice and performed proteomic analysis. The common pathways affected in these two different mouse models would be likely to be *bona fide* one involved in the PD pathogenesis. Before the proteome analysis of the model mice, neurological pathologies of those mice were carefully accessed. Mouse brains were harvested at 6 months after post-injection. For more accurate protein quantification of mouse samples, stable isotope labeling with amino acid in mammal (SILAM) mice were generated by labeling with heavy lysine and spiked into the ventral midbrain samples from PD model or control mice. The proteins were extracted from those samples followed by Lys-C digestion. The relative protein abundances were measured by LC-MS/MS after fractionating the peptides by high pH RPLC (Fig. 1).

**Figure 1.**
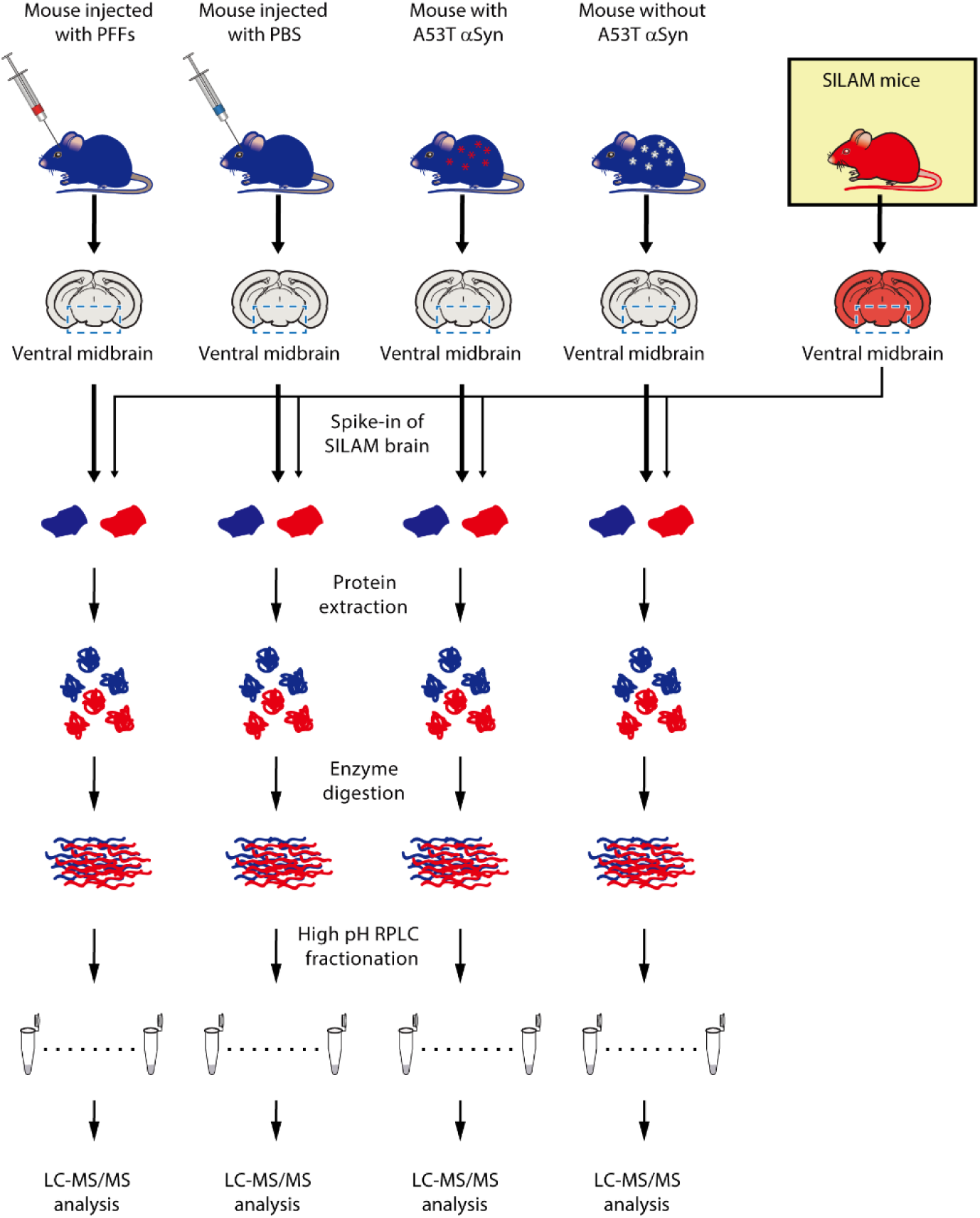
Schematic diagram of the proteome analyses of the ventral midbrain of intrastriatal α-synuclein PFF mice and AAV1-tTA/TetP-hA53T-α-synuclein Tg mice. Brains from the two PD mouse models were subjected to quantitative proteome analysis. For the accurate protein abundance quantification, mice that were labeled with heavy Lys for three generations were spiked in before lysis of the brain samples. The ventral midbrain regions from each mouse were harvested and the SILAM brains were added to the PD model and their control mice. After protein extraction from the ventral midbrains, the proteins were digested with Lys-C and trypsin subsequently. The peptides were fractionated by high pH RPLC followed by LC-MS/MS analysis.

### The generation and validation of neurological pathologies of α-synuclein PFFs-induced mouse model of PD

To dissect pathways implicated in the pathogenesis of α-synuclein PFFs-induced mouse model of PD, the model was generated by injecting either PFF or PBS to the striatum via stereotactically unilateral injection as previously reported (8, 19, 22, 23, 30) (Fig. 2A). Before starting the proteome analysis of the ventral midbrain region of the mice, validation of the neuropathological symptoms was crucial. To this end, the assessment of DA neuronal death and behavioral tests of mouse models were conducted. The DA neuronal degeneration of α-synuclein PFFs-induced mouse was assessed by TH-immunoreactivity and Nissl staining using unbiased stereological counting. After 6 months of post-injection, about 62% of the TH positive neurons and 59% of Nissl-positive neurons were degenerated in the substantia nigra (SN) region of α-synuclein PFFs injected hemisphere (Fig. 2B and C). The amount of TH-positive fibers in the striatum was also reduced after 6 months of α-synuclein PFFs injection (Fig. 2D). Behavioral deficit was accessed by the rotarod and pole tests. α-Synuclein PFFs injection significantly led to behavioral deficits in the rotarod and pole tests (Fig. 2E). The p-α-synuclein pathologies and propagation were also detected in the various regions including the SN after 6 months of α-synuclein PFFs injection (Fig. 2F and G) as previously described (8, 19, 22, 23, 30).

**Figure 2.**
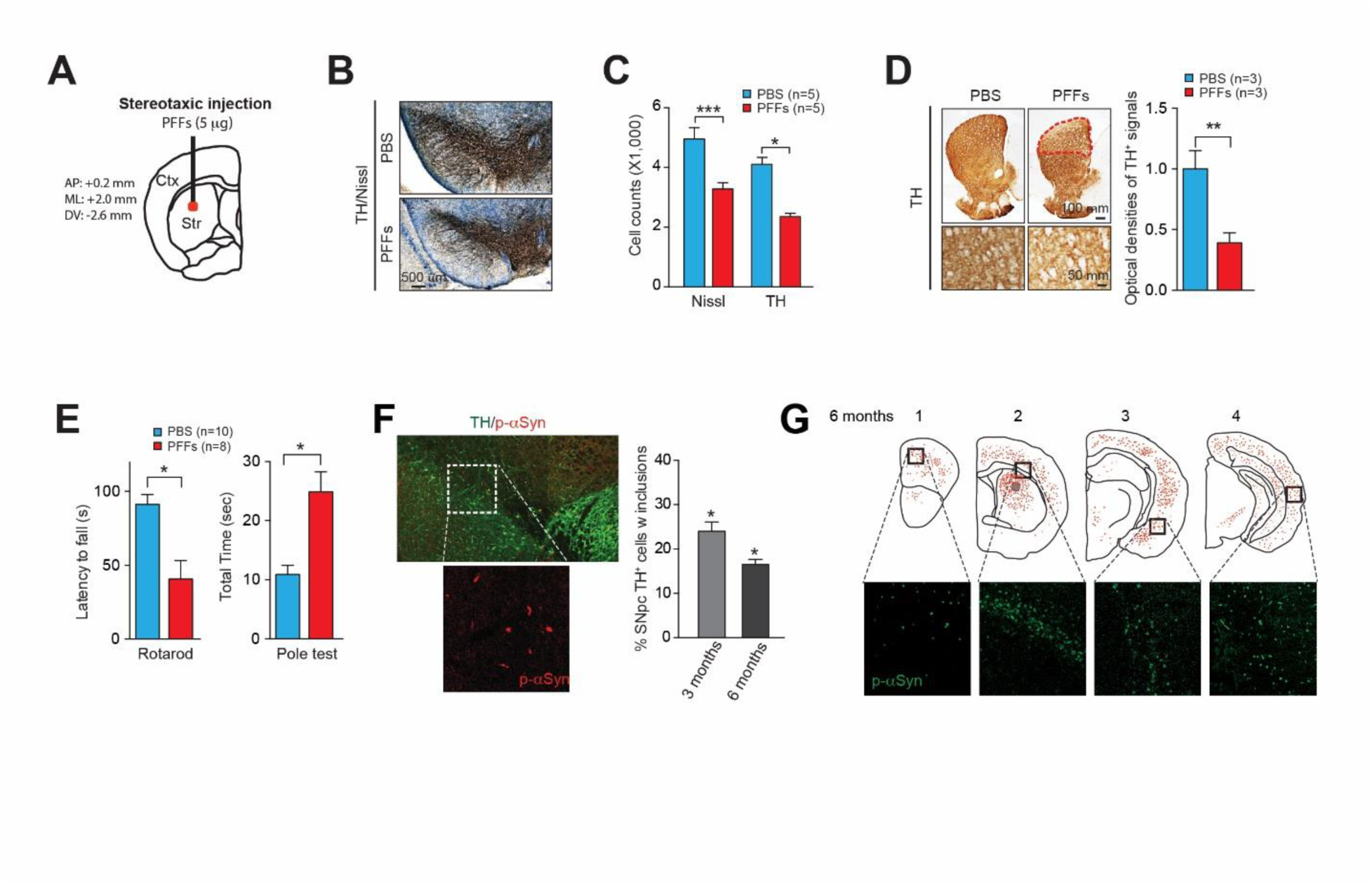
The generation and characterization of intrastriatal α-synuclein PFF injection mouse model of PD. **A,** Schematic illustration of the coordination for α-synuclein PFF (5 μg) stereotaxic injection. **B,** Representative TH-immunostaining images in the SN of α-synuclein PFF or PBS-injected mice. **C,** Stereological counting of the number of Nissl- and TH-positive neurons in the SN after 6 months of PBS or α-synuclein PFF injection (n=5 per group). **D,** Representative TH-immunostaining images in the striatum of PBS or α-synuclein PFF injected mice. The TH-immunopositive fiber densities were quantified by using ImageJ software (n=5 per group). **E,** Assessments of the behavioral deficits measured by rotarod test and pole test. **F,** Representative p-α-synuclein immunostaining images in the SN of PBS or α-synuclein PFF injected mice. The formation of pS129-α-synuclein positive inclusion in TH-positive cells were quantified and represented as a graph (n=5, each group). **G,** Distribution of LB/LN-like pathology in the CNS of α-synuclein PFFs-injected hemisphere. Student’s t test was used to test for the statistical analysis. *\*P* < 0.05, *\*\*P* < 0.01, *\*\*\*P* < 0.001.

### The generation and validation of neurological pathologies of AAV1-tTA injected TetP-hA53T α-synuclein transgenic mouse model

To identify pathways involved in the pathogenesis of AAV1-tTA injected TetP-hA53T α-synuclein transgenic (Tg) mouse model, the model mice were generated by injecting either AAV1-tTA virus or AAV1-GFP virus into the ventral midbrain region of conditional TetP-hA53T α-synuclein mice (18) via stereotactically unilateral injection (Fig. 3A). Before starting proteome analysis of the ventral midbrain regions of the mice, validation of the neuropathological symptoms was conducted. After 6 months of intranigral AAV1-tTA virus injection, 2.3-fold increase of α-synuclein was observed in the ventral midbrain of AAV1-tTA/TetP-hA53T α-synuclein Tg mouse leading to a reduction in TH protein levels (Fig. 3A-C). The hA53T α-synuclein overexpression induced a significant loss of TH and Nissl positive neurons in the SNpc (Fig. 3D and E). Moreover, striatal TH immunoreactivity was reduced by overexpression of hA53T α-synuclein (Fig. 3F). hA53T α-synuclein overexpression also significantly elicited the behavioral deficits as measured by the accelerating rotarod test and the pole test (Fig. 3G).

**Figure 3.**
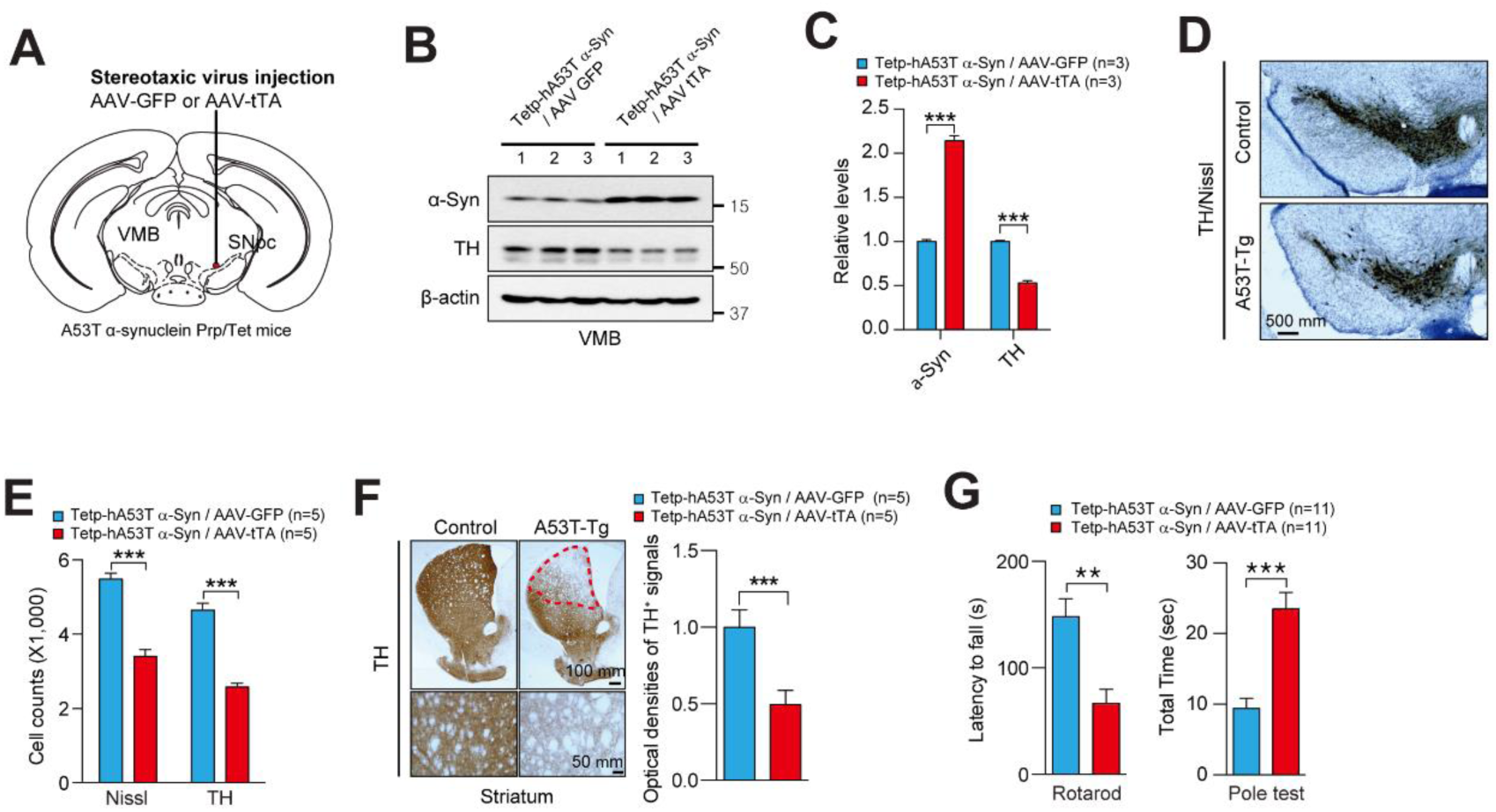
The generation and characterization of AAV1-tTA/TetP-hA53T-α-synuclein Tg mouse model of PD. **A**, Schematic of experiments for mice with stereotactic injection of AAV1-tTA. **B,** Representative immunoblots of TH, α-synuclein, and β-actin in the ventral midbrain of hA53T α-synuclein Tg mice. **C**, Quantification of α-synuclein, and TH protein levels in the ventral midbrain of hA53T α-synuclein mice normalized to β-actin. **D**, Representative TH immunostaining of midbrain sections from the SN of AAV1-tTA-injected TetP-hA53T α-synuclein mice and AAV1-GFP-injected TetP-hA53T α-synuclein mice. **E**, Stereological assessments of TH and Nissl positive neurons in the SN of AAV1-tTA-injected TetP-hA53T α-synuclein mice and AAV1-GFP-injected TetP-hA53T α-synuclein mice. (n = 5 per group). **F**, Representative TH immunostaining of mouse striatal sections from AAV1-tTA-injected TetP-hA53T α-synuclein mice and AAV1-GFP-injected TetP-hA53T α-synuclein mice at 6-month post injections. Quantifications of dopaminergic fiber densities in the striatum by using Image J software. (n = 5 per group). **G,** Assessments of the behavioral deficits measured by rotarod test and pole test. Student’s t test was used to test for the statistical analysis. *\*\*P* < 0.01, *\*\*\*P* < 0.001 vs. α TetP-hA53T α-synuclein mice with AAV1-GFP.

### Proteomic analysis of the ventral midbrain from intrastriatal α-synuclein PFF injection mice and AAV1-tTA/TetP-hA53T-α-synuclein Tg mice

To investigate the common pathways affected in the ventral midbrain of two different mouse models of PD, we performed the proteome analysis with ventral midbrain samples prepared from the intrastriatal α-synuclein PFF injection mice and AAV1-tTA/TetP-hA53T-α-synuclein Tg mice (Fig. 1). The ventral midbrain samples were subject to proteome analysis by extracting proteins, enzyme digestion, pre-fractionation, and mass spectrometry analysis. For more accurate protein quantification, SILAM mice were generated and spiked into the mouse samples for quantification before protein extraction. Before spiking in the ventral midbrain samples labeled with heavy Lys, the labeling efficiency of the proteins in the serum and brain were measured. The ratio of heavy to light was found to be approximate >99%, and a few of the randomly selected targets were manually verified by scrutinizing the MS1 and the representative MS/MS spectra for the peptides AEGAGTEEEGTPK (Brain acid soluble protein), STEFNEHELK (Visinin like proteins) from the brain tissue and TSDQIHFFFAK (Antithrombin III) and HITSLEVIK (Platelet factor IV) from the serum sample (Supplemental Fig. S1A-D). The peptide samples were injected twice to the mass spectrometry. Approximately 6,800 and 7,200 proteins were identified from intrastriatal α-synuclein PFF injection mice and AAV1-tTA/TetP-hA53T-α-synuclein Tg mice, respectively. When the proteins from intrastriatal α-synuclein PFF injection mice and AAV1-tTA/TetP-hA53T-α-synuclein transgenic mice were combined, in total, 8,355 proteins were identified, and 5,647 proteins were identified in common (Supplemental Fig. S2). When statistical analyses were performed, 724 and 1581 proteins were differentially regulated by α-synuclein PFF injection and nigral hA53T-α-synuclein overexpression, respectively (Fig. 4A and B). Among them, 149 proteins were differentially regulated in common. Importantly, TH protein, one of the protein specific to dopaminergic neurons, showed 3.3- and 3.4- fold decrease in α-synuclein PFF injection and nigral hA53T-α-synuclein overexpression, respectively. These results are consistent with the assessments of the dopaminergic neuronal death shown in Figure 1 and 2. These results also suggest that the quantitative proteomic experiments were properly conducted. In the intrastriatal α-synuclein PFF injection mice, collagen type VI alpha 3 chain (COL6A3) showed the highest upregulation with ∼60 folds followed by collagen type XII alpha chain (COL12A) and Transgelin (TAGLN) with statistical significance. In AAV1-tTA/TetP-hA53T-α-synuclein transgenic mice, Serpin peptidase inhibitor clade A (SERPINA1A) showed the highest upregulation with ∼20 folds of increase followed by immunity-related GTPase family M member 1(IRGM1) and ATPase sarcoplasmic/endoplasmic reticulum Ca^2+^ transporting 2 (ATP2A2). In both mouse models, the level of downregulation was mostly less than 4 folds except for myelin protein zero (MPZ) with ∼60 folds downregulation in the AAV1-tTA/TetP-hA53T-α-synuclein transgenic mice. These results suggest that the dopaminergic neurons in the brains of α-synuclein PFF injection and nigral hA53T-α-synuclein overexpression PD mouse models were damaged and the quantitative proteomic measurement of protein abundance was correctly detected this change.

**Figure 4.**
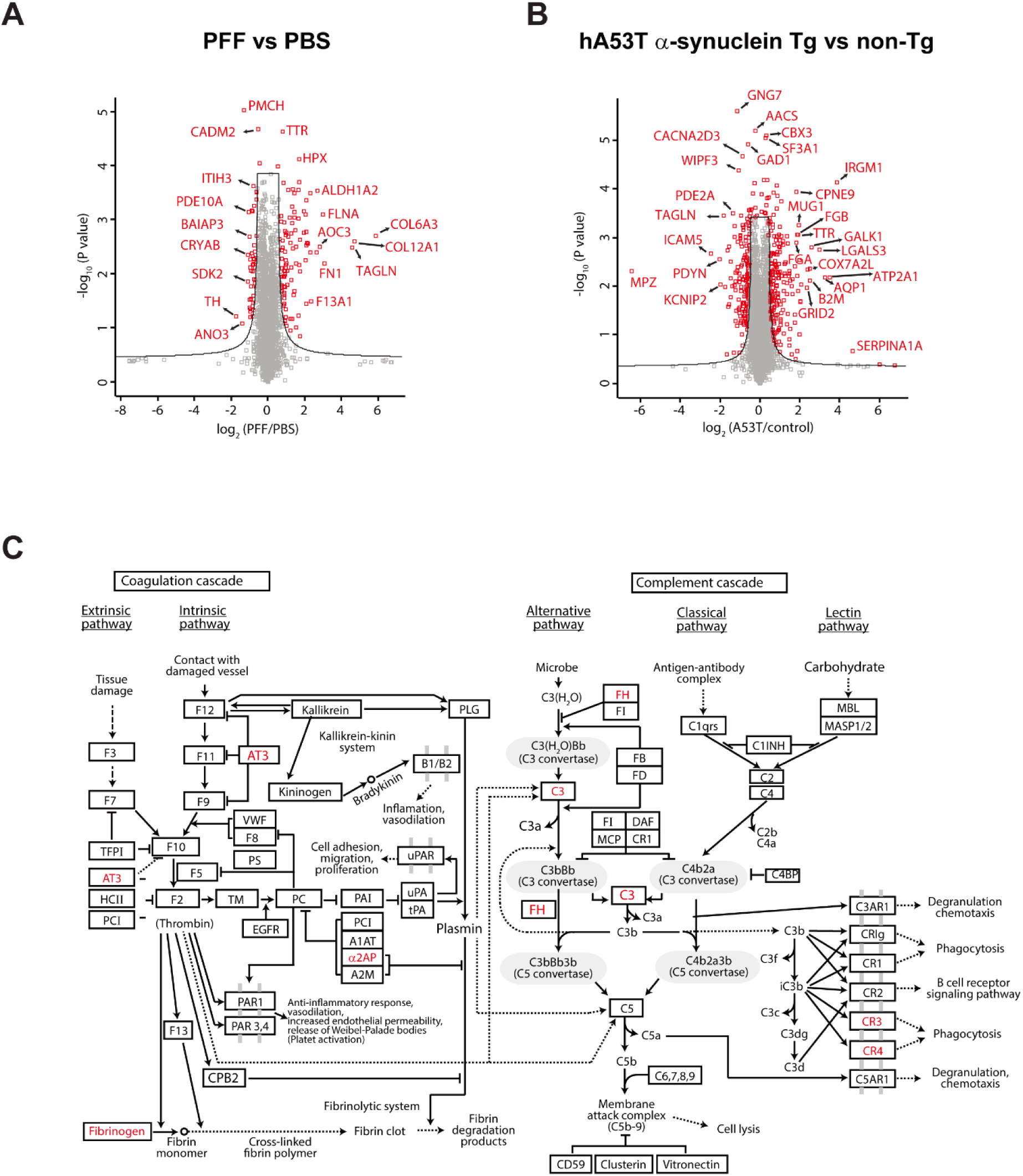
Volcano plot and pathway analyses of quantified proteins in the two mouse models of PD. **A,** The proteins quantified in PFF injected and PBS injected mice were analyzed for differentially expressed ones in PFF injected mice. The cutoff used to select differential proteins was q-value < 0.05. **B,** The proteins quantified in hA53T α-synuclein Tg and control mice were analyzed for differentially expressed ones in hA53T α-synuclein Tg mouse. The cutoff used to select differential proteins was q-value <0.05. **C,** The proteins differentially expressed in both PFF injected mice and A53T α-synuclein Tg mice were analyzed for enriched pathways. Complement cascade and coagulation cascade were most enriched one for the differential expressed proteins both in PFFs injected mice and A53T α-synuclein Tg mice. The pathway analysis was performed in KEGG incorporated in DAVID.

**Figure 5.**
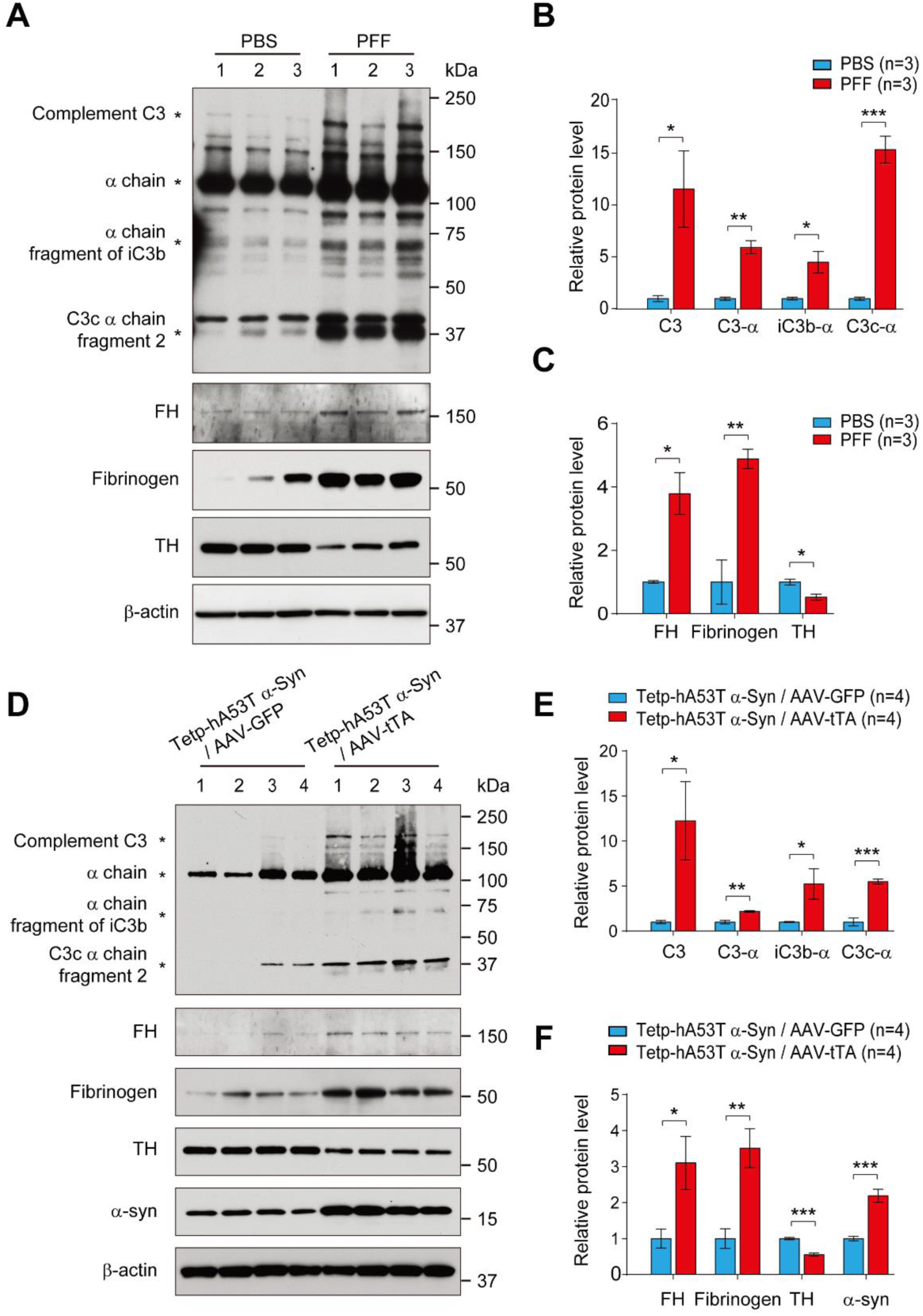
Western blot validation of specific proteins identified in the proteomic analysis and involved in coagulation cascade and component cascade. **A,** Representative immunoblots of complement C3, alpha chain of C3, alpha chain fragment of iC3b, and C3c alpha chain fragment 2, Factor H (FH), Fibrinogen, TH and β-actin in the ventral midbrain of α-synuclein PFF injected mice. **B,** Quantification of complement C3, alpha chain of C3, alpha chain fragment of iC3b, and C3c alpha chain fragment 2 protein levels in the ventral midbrain normalized to β-actin. **C,** Quantification of FH, Fibrinogen, and TH protein levels in the ventral midbrain normalized to β-actin. Error bars represent the mean ± S.E.M (n=3 mice per groups). Student’s t test for statistical significance. *\*P* < 0.05, *\*\*P* < 0.01, *\*\*\*P* < 0.001. D, Representative immunoblots of complement C3, alpha chain of C3, alpha chain fragment of iC3b, and C3c alpha chain fragment 2, Factor H (FH), Fibrinogen, TH and β-actin in the ventral midbrain of hA53T α-synuclein Tg mice. **E,** Quantification of complement C3, alpha chain of C3, alpha chain fragment of iC3b, and C3c alpha chain fragment 2 protein levels in the ventral midbrain normalized to β-actin. **F,** Quantification of FH, Fibrinogen, TH, α-synuclein protein levels in the ventral midbrain normalized to β-actin. Error bars represent the mean ± S.E.M (n=3 mice per groups). Student’s t test for statistical significance. *\*P* < 0.05, *\*\*P* < 0.01, *\*\*\*P* < 0.001.

### Pathway analysis of the differentially expressed proteins common in intrastriatal PFFs and AAV1-tTA/TetP-A53T Tg mouse models of PD

Next, using data that emerged from the proteome analysis of two different types of mouse model of PD, we performed the pathway analysis on DAVID in order to investigate the signaling pathways commonly affected in the models. Since the two different model systems were designed to develop the toxicity to dopaminergic neurons by fostering α-synuclein aggregation, we reasoned that those commonly differentially regulated proteins in the two model systems would have a much higher chance to play a role in PD pathogenesis. Thus, we performed the pathway analysis with differentially expressed proteins in two PD mouse model systems giving 26 enriched pathways (Table 2). Complement and coagulation cascades were identified as the most enriched pathways (Fig. 4C) followed by spliceosome, dopaminergic synapse, cocaine addiction, and platelet activation. Intriguingly, the dopaminergic synapse was the third most enriched pathway suggesting that the dopaminergic pathway was altered by α-synuclein PFF injection and nigral hA53T-α-synuclein overexpression.

**Table 2.** Enriched pathways by the differential proteins common in both α-synuclein PFFs injected and hA53T α-synuclein Tg mice

| Term | Count | P Value | Genes |
| --- | --- | --- | --- |
| Complement and coagulation cascades | 7 | 1.17E-04 | *FGG, FGA, SERPINF2, C3, FGB, SERPINC1, CFH* |
| Spliceosome | 8 | 3.97E-04 | *U2SURP, DHX15, SNW1, HNRNPC, SNRPC, HNRNPU, RBM25, SF3B3* |
| Dopaminergic synapse | 8 | 4.16E-04 | *DDC, PPP1R1B, GNAI1, ADCY5, TH, PLCB1, PPP2R2C, PPP2R2A* |
| Cocaine addiction | 5 | 0.001517695 | *DDC, PPP1R1B, GNAI1, ADCY5, TH* |
| Platelet activation | 7 | 0.00214858 | *FGG, TLN1, FGA, FGB, GNAI1, ADCY5, PLCB1* |
| Amphetamine addiction | 5 | 0.004779233 | *DDC, PPP1R1B, ADCY5, TH, PPP3R1* |
| Renin secretion | 5 | 0.005875053 | *PDE1B, GNAI1, ADCY5, PPP3R1, PLCB1* |
| Tight junction | 6 | 0.01368547 | *MPDZ, GNAI1, MYH9, CTNNA1, PPP2R2C, PPP2R2A* |
| Estrogen signaling pathway | 5 | 0.017765421 | *HSP90B1, HSP90AA1, GNAI1, ADCY5, PLCB1* |
| Focal adhesion | 7 | 0.018970256 | *TLN1, ITGA6, DIAPH1, TNC, FLNC, LAMB1, FN1* |
| Pathways in cancer | 10 | 0.019510174 | *HSP90B1, HSP90AA1, ITGA6, GNAI1, ADCY5, STAT1, PLCB1, LAMB1, CTNNA1, FN1* |
| Chagas disease | 5 | 0.020931721 | *C3, GNAI1, PLCB1, PPP2R2C, PPP2R2A* |
| Gastric acid secretion | 4 | 0.03715889 | *EZR, GNAI1, ADCY5, PLCB1* |
| Serotonergic synapse | 5 | 0.045921293 | *DDC, GNAI1, ADCY5, PLCB1, TPH2* |
| Alcoholism | 6 | 0.055493563 | *DDC, PPP1R1B, GNAI1, ADCY5, H2AFY2, TH* |
| Gap junction | 4 | 0.057672934 | *GNAI1, ADCY5, TUBA4A, PLCB1* |
| GABAergic synapse | 4 | 0.059307468 | *PLCL1, GNAI1, ADCY5, GAD1* |
| ECM-receptor interaction | 4 | 0.060963804 | *ITGA6, TNC, LAMB1, FN1* |
| PI3K-Akt signaling pathway | 8 | 0.066992193 | *HSP90B1, HSP90AA1, ITGA6, TNC, LAMB1, PPP2R2C, FN1, PPP2R2A* |
| Adrenergic signaling in cardiomyocytes | 5 | 0.067254891 | *GNAI1, ADCY5, PLCB1, PPP2R2C, PPP2R2A* |
| Regulation of actin cytoskeleton | 6 | 0.067675961 | *EZR, ITGA6, DIAPH1, MSN, IQGAP1, FN1* |
| Morphine addiction | 4 | 0.069566791 | *PDE1B, PDE2A, GNAI1, ADCY5* |
| Tryptophan metabolism | 3 | 0.083203379 | *DDC, CAT, TPH2* |
| Staphylococcus aureus infection | 3 | 0.092484044 | *FGG, C3, CFH* |
| Protein processing in endoplasmic reticulum | 5 | 0.092914276 | *HSPH1, HSP90B1, HSP90AA1, HSPA4L, UBE2E2* |
| cGMP-PKG signaling pathway | 5 | 0.097590751 | *PDE2A, GNAI1, ADCY5, PPP3R1, PLCB1* |

### Validation of specific proteins identified in the proteomic analysis

To further validate the accuracy of the proteomic analysis, the levels of several target proteins were assessed by Western blot analysis with the ventral midbrain samples prepared from the intrastriatal α-synuclein PFF injection mice and AAV1-tTA/TetP-hA53T-α-synuclein Tg mice. The intrastriatal α-synuclein PFFs injection significantly increased the protein levels of complement C3, alpha chain, alpha chain fragment of iC3b, and C3c alpha chain fragment 2 (Fig. 6A and B) as well as Factor H (FH) (Fig. 6C) that are involved in the complement cascade in the ventral midbrain with a reduction in TH protein levels (Fig. 6C). In addition, the protein level of fibrinogen in coagulation cascade was markedly increased in the ventral midbrain of intrastriatal α-synuclein PFFs PD mouse model. Similar results were obtained in AAV1-tTA/TetP-hA53T-α-synuclein Tg mice. Nigral hA53T-α-synuclein overexpression led to an increase in the protein levels of complement C3, alpha chain, alpha chain fragment of iC3b, and C3c alpha chain fragment 2 (Fig. 6D and E) as well as FH (Fig. 6F), all of which are involved in the complement cascade in the ventral midbrain with reduction in TH protein levels (Fig. 6F). In addition, the protein level of fibrinogen in coagulation cascade was markedly increased in the ventral midbrain of the AAV1-tTA/TetP-hA53T-α-synuclein Tg mice (Fig. 6F).

## Discussion

In this study, we conducted the proteome analysis of the ventral midbrain from two sophisticated mouse models of PD: α-synuclein PFFs injection mouse model that represents sporadic PD (8) and AAV1-tTA/TetP-hA53T α-synuclein transgenic (Tg) mouse model that represents familial PD (18), to investigate pathways inextricably involved in the α-synuclein induced dopaminergic neurodegeneration. The recognizable finding from the proteome analysis is the observation that coagulation cascade and complement cascade are potentially involved in the neurodegeneration progress in PD.

Mass spectrometry-based proteomics technologies have been widely used for the identification of proteins that are differentially expressed in disease animal models on a global scale (12–15). To identify differentially expressed proteins, accurate quantifications of proteins are essential. Currently, multiple protein quantification approaches such as label-free quantification, metabolic labeling, isobaric mass tags are available (31). Among them, the metabolic labeling-based protein quantification such as SILAM provides the most accurate protein quantification since the experimental variation is minimized by mixing brain samples with brain samples labeled with heavy an amino acid before starting a protein extraction. In this study, we quantified protein abundances by adding SILAM-labeled mouse brain samples and accurately quantified protein abundance.

When the pathway analyses were performed separately, the enriched pathways in two different PD mouse model systems were quite different (Supplemental Table S3). On the other hand, when only differential proteins in both PD mouse models were counted, dopaminergic synapse pathway was shown up at top 3 suggesting that the pathway by the differential proteins in common were leading us to the right affected pathway. Strikingly, the complement pathway and coagulation cascade pathway were the most enriched (Fig. 3C).

The complement system is well-known as an essential branch of the innate immune system. Complement activation is elevated in human PD brain, especially in the presence of Lewy bodies (LB). Yamada and colleagues reported intra-and extra-neuronal LB in the substantia nigra of patients with PD were stained with both antibodies for early-stage (C3d and C4d) and late-stage (C7 and C9) complement proteins (32). C3d and C4d staining for the LB were continuously reported in the brainstem of patients with dementia with Lewy bodies (DLB) (33). Over time, Loeffler and colleagues (34) reported a significantly higher proportion of iC3b^+^ neurons in normal aged-specimens than in normal young specimens and a more considerable increase in PD specimens. Interestingly, they also reported that the percentage of iC3b^+^ neurons in the PD specimens was significantly higher than in the Alzheimer’s disease (AD) specimens.

In the same context, complement C3 plays a role in the complement cascade as a central molecule that is closely related to the central nervous system (CNS) synapse elimination and brain aging (35, 36). The region-specific synapse and neuron loss as well as cognitive decline caused by normal aging have been reported to be spared in C3 knockout mice (36). Recently, it has been well documented that complement C3 and its downstream iC3b/CR3 signaling play an important role in AD. Shi and colleagues found that C3 deficiency in amyloid precursor protein/presenilin 1 (APP / PS1) mice confers neuroprotection against age- and AD-related neurodegeneration and cognitive decline, and C3 has been shown to be detrimental to synapse and neuronal function in plaque-rich aged brains (37).

In addition, growing evidence suggest that complementary C3 contributes to pathogenesis in neurodegenerative diseases. It is important to note that A1 astrocytes identified in post-mortem tissue from individuals with AD, PD, Multiple sclerosis, Amyotrophic lateral sclerosis, and Huntington’s disease were C3-positive (38). Moreover, consistent with our finding, C3d immunoreactivity in the ventral midbrain with robust loss of DA neuron was induced by intrastriatally injected α-synuclein PFF (23) suggesting that complementary pathway may contribute to drive neurodegeneration in two mouse models of PD. The brains in the two mouse models of PD were likely to include full of A1 astrocytes releasing of complement factor C3, which may contribute to pathological α-synuclein LB pathology and neurodegeneration in the two mouse models of PD. Further investigation is necessary to understand the role of the complement system in the pathogenesis of PD.

Factor H (FH), the main regulator of this pathway, prevents formation and promotes the dissociation of C3 convertase enzyme (39, 40). Several studies have suggested cerebrospinal fluid (CSF) C3 and FH levels could serve as biomarkers for AD and PD (41–44). According to the alteration of FH expression in neurodegeneration, the expression level of FH was significantly increased in the postmortem brain of AD cases than in the normal elderly (ND) cases (45). Importantly, immunohistochemical study of FH showed intense immune responses throughout the AD brain, including aβ plaques, neurons, and glia, while ND did not (45). Over time, the FH protein has been demonstrated as a potential biomarker for AD (46, 47). Strohmeyer and colleagues (45) demonstrated possible interaction of FH with aggregated fibrillar Aβ but not nonfibrillar Aβ.

Under normal conditions, the complement system plays a number of vital roles in physiological process during brain development and homeostasis (48). At the disease conditions, complement synthesis in brain cells is increasing significantly in parallel with tissue damage (49). During the early disease onset, abnormal upregulation and deposition of complement contributes to synapse loss and cognitive dysfunction (50, 51). In addition, there is increasing evidence that inhibiting the complement pathway is effective in improving the effects of neurodegenerative pathologies (37, 52). In this study, we showed that complement components including C3, its fragments and FH were significantly induced in ventral midbrain (VMB) of the two PD mouse models. The finding suggests that the complement pathway probably mediates neurodegeneration by α-synuclein, such as neuronal loss. C3-knockout (KO)/APP-transgenic mice and C3ar1-KO/PS19-transgenic mice also exhibit reduced microglial lysosomal content and astrocyte GFAP levels (37, 53). And the main cell types of brain parenchyma, such as astrocytes and microglia, can take up extracellular α-synuclein aggregates and degrade them (54). It has been reported that microglial phagocytosis is one of the major causes of complement-mediated synapse loss (50). This suggests that the complement system modulates immune cell function and thereby contributes to neurodegeneration. On the other hand, microglia express receptors for various complement components, including C1q and C3 cleavage products, and thus the complement system plays a crucial role in microglial activation (55, 56). Taken together, the complement system maybe contributes to maintaining homeostasis by coordinating immune functions in the healthy state or early stages of PD. However, as the PD deteriorates to the late stage, an abnormal increase in the amount of complement components and activation of the complement system will lead to an imbalance of homeostasis, which will become detrimental and eventually lead to neurodegeneration.

The blood coagulation factor fibrinogen is released into CNS parenchyma after blood–brain barrier (BBB) breakdown and converted to insoluble fibrin, a major proinflammatory matrix that activates the innate immune response (57, 58). An emerging study using genetic or pharmacological depletion of fibrinogen indicates that fibrinogen is not only an indicator of BBB destruction, but it also plays a role in various animal models of neurological disease, particularly in multiple sclerosis (59), AD (60), brain trauma (61), and nerve injury (62). According to a large population-based cohort study, high fibrinogen levels were associated with higher prevalence and incidence of PD in participants aged 76 years and older (63). In addition, fibrinogen showed a similar 9.4-fold extravascular increase in the post-commissural putamen of PD patients (64). In fact, fibrinogen deposition increases simultaneously with the increase in amyloid-β in the brain as AD pathology progresses (60, 65). In addition, increased fibrinogen levels are notedly associated with an increased risk of dementia and AD (66, 67).

In summary, our results suggest that coagulation cascade and complement cascade potentially contribute to alpha-synuclein-induced neurodegeneration in PD. These pathways may provide a meaningful window into new drug targets or neuroprotein therapeutic strategies to treat PD

## Abbreviations

PFF: Pre-formed fibrils
PD: Parkinson’s disease
AD: Alzheimer’s disease
LB: Lewy bodies
SILAM: Stable isotope labeling with amino acid in mammal
TH: Tyrosine hydroxylase
DA: Dopamine
SNpc: Substantia nigra par compacta
MPTP: 1-methyl-4-phenyl-1,2,3,6-tetrahydropyridine
6-OHDA: 6-Hydroxydopamine
tTA: Tetracycline-regulated transactivator
TetP: Tetracycline promoter
AAV: Adeno-associated virus
LC-MS/M: Liquid Chromatography with tandem mass spectrometry
Tg: Transgenic
CNS: Central nervous system

## Footnotes

The authors acknowledge the joint participation of the Adrienne Helis Malvin Medical Research Foundation and the Diana Helis Henry Medical Research Foundation through its direct engagement in the continuous active conduct of medical research in conjunction with The Johns Hopkins Hospital and the Johns Hopkins University School of Medicine and the Foundation’s Parkinson’s Disease Program M-2014, H-1, H-2013.

## Author contributions

C.H.N., D.K, V.L.D., T.M.D, A.P. and H.S.K. designed research; C.H.N., S.X.M., D.K., S.P.Y., Y.X., O.P., and R.S.N. conducted the experiment; C.H.N. S.X.M., performed data analysis. C.H.N, S.X.M., D.K, S.P.Y., J.T., T.M.D., A.P and H.S.K. wrote manuscript; T.M.D., A.P. and H.S.K. supervised research.

---

**Supplemental Table S1.** Antibodies used in this study

| Antibodies | Source/Cat. No./Ref. | Host | Dilution |
| --- | --- | --- | --- |
| α-Synuclein | BD Bioscience (610787) | Mouse | 1:1,000 (WB) |
| p-α-synuclein<sup>Ser129</sup> | Biolegend (825701) | Mouse | 1:1,000 (ICH, IF) |
| Tyrosine Hydroxylase (TH) | Novus Biologicals<br>(NB300-19) | Rabbit | 1:1,0000 (WB)<br>1:1,000 (IHC, IF) |
| Factor H | Abcam (ab8842) | Sheep | 1:1,000 (WB) |
| C3 | Abcam (ab200999) | Rabbit | 1:1,000 (WB) |
| Fibrinogen | Abcam (ab189490) | Rabbit | 1:1,000 (WB) |
| β-actin-HRP | Sigma-Aldrich (A3854) | | 1:10,000 (WB) |

Supplemental Table S2. List of entire identified and quantified proteins

Supplemental Table S3. List of enriched pathways in α-synuclein PFF injected mice and hA53T α-synuclein Tg mice.

**Supplemental figure S1.**
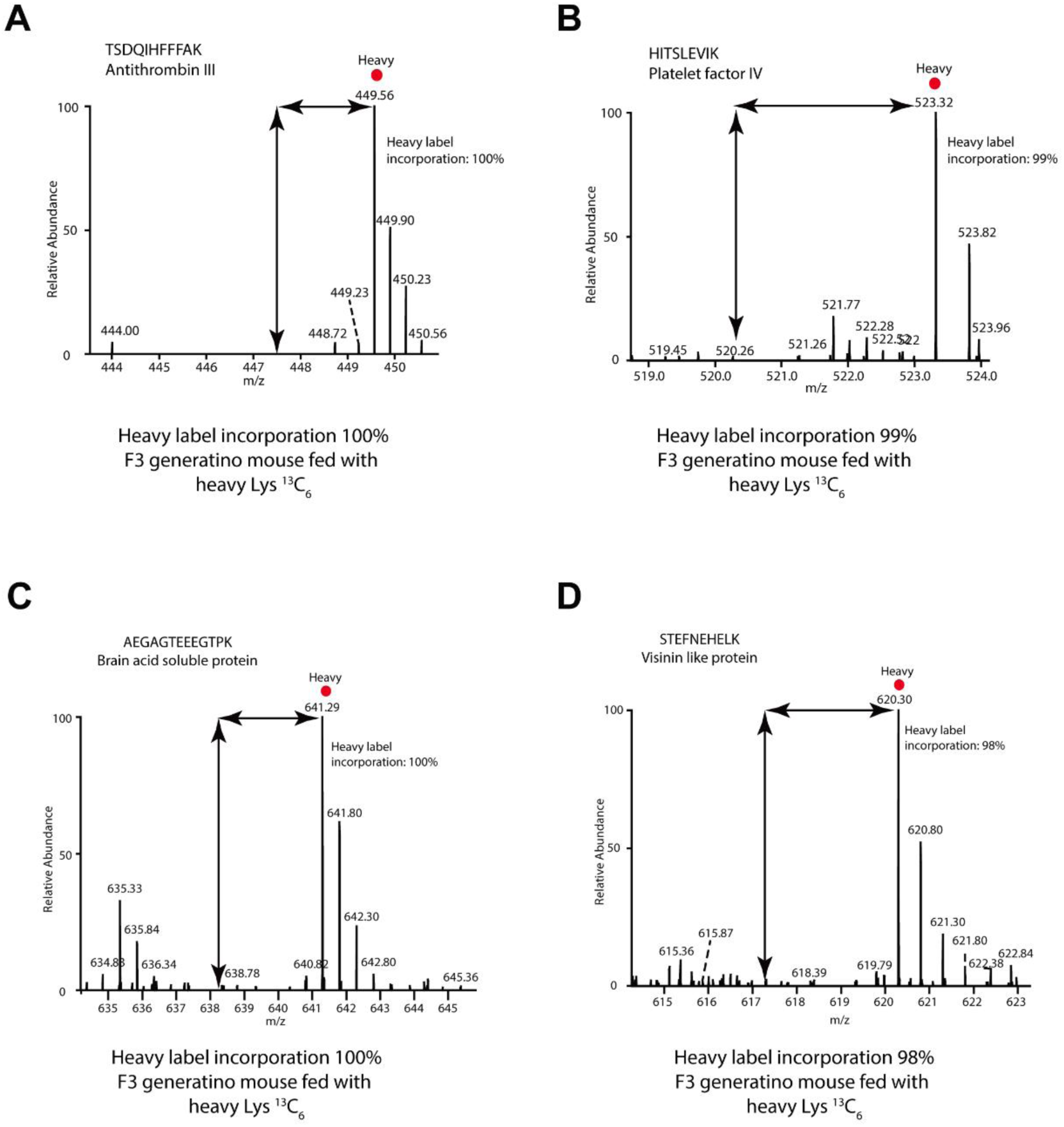
The labeling efficiency of serum and brain proteins from the SILAM mice. **A**, The heavy to light ratio of a peptide, TSDQIHFFFAK derived from a serum protein, antithrombin III. **B**, The heavy to light ratio of a peptide, HITSLEVIK derived from a serum protein, platelet factor IV. **C**, The heavy to light ratio of a peptide, AEGAGTEEEGTPK derived from a brain protein, brain acid soluble protein. **D**, The heavy to light ratio of a peptide, STEFNEHELK derived from a brain protein, visinin like protein.

**Supplemental figure S2.**
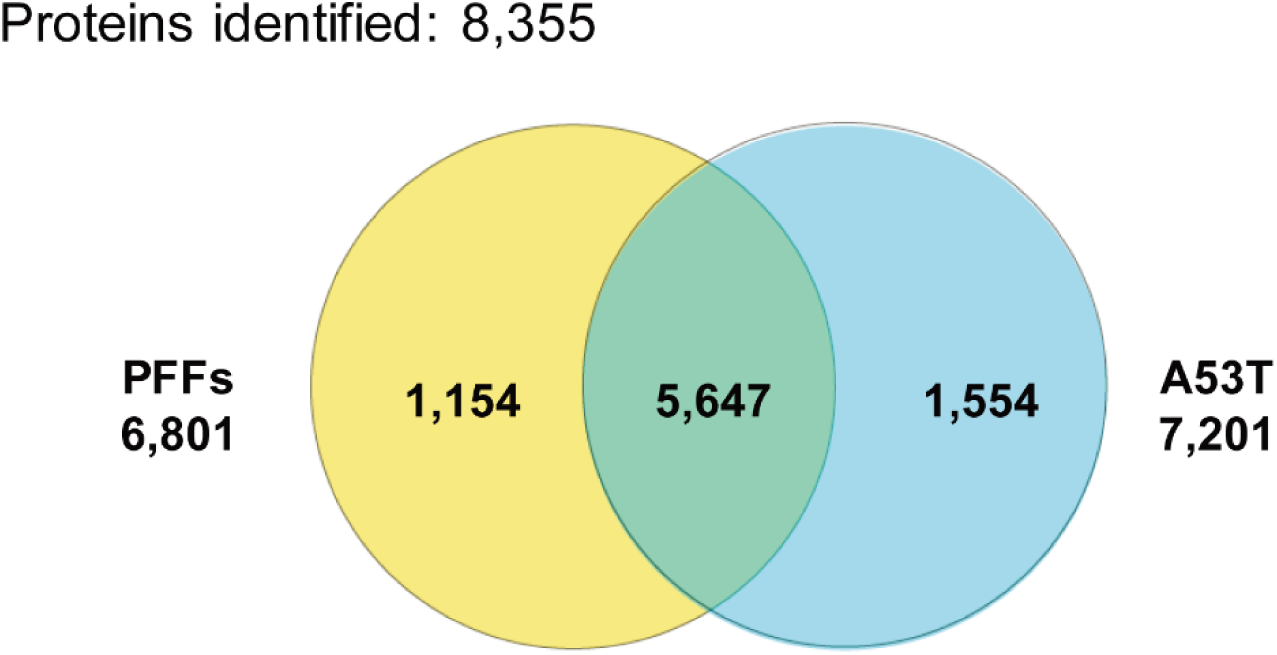
The number of proteins identified from the PD mouse model with intrastriatal α-synuclein PFF injection (left) and the PD mouse model with human A53T-α-synuclein expression (right).

